# Increased prostaglandin-D_2_ in male but not female STAT3-deficient hearts shifts cardiac progenitor cells from endothelial to white adipocyte differentiation

**DOI:** 10.1101/2020.04.24.059287

**Authors:** Elisabeth Stelling, Melanie Ricke-Hoch, Sergej Erschow, Steve Hoffmann, Anke Katharina Bergmann, Maren Heimerl, Stefan Pietzsch, Karin Battmer, Alexandra Haase, Britta Stapel, Michaela Scherr, Jean-Luc Balligand, Ofer Binah, Denise Hilfiker-Kleiner

**Author notes:** Corresponding author: (DH-K). These authors contributed equally to this work (Elisabeth Stelling, Melanie Ricke-Hoch).

## Abstract

Cardiac levels of the signal transducer and activator of transcription factor-3 (STAT3) decline with age, and male but not female mice with a cardiomyocyte-specific STAT3 deficiency (CKO) display premature age-related heart failure associated with reduced cardiac capillary density. In the present study isolated male and female CKO-cardiomyocytes exhibit increased prostaglandin (PG)-generating cyclooxygenase-2 (COX-2) expression. The PG-degrading hydroxyprostaglandin-dehydrogenase-15 (HPGD) expression is only reduced in male cardiomyocytes, which is associated with increased PGD_2_ secretion from isolated male but not female CKO-cardiomyocytes. Reduced HPGD expression in male cardiomyocytes derive from impaired androgen-receptor-(AR)-signaling due to loss of its co-factor STAT3. Elevated PGD_2_ secretion in males is associated with increased white adipocyte accumulation in aged male but not female hearts. Adipocyte differentiation is enhanced in isolated SCA-1^+^-cardiac-progenitor-cells (CPC) from young male CKO-mice compared to the adipocyte differentiation of male wildtype (WT)-CPC and CPC isolated from female mice. Epigenetic analysis in freshly isolated male CKO-CPC display hypermethylation in pro-angiogenic genes (*Fgfr2*, *Epas1*) and hypomethylation in the white adipocyte differentiation gene *Zfp423* associated with upregulated ZFP423 expression and a shift from endothelial to white adipocyte differentiation compared to WT-CPC. The expression of the histone-methyltransferase EZH2 is reduced in male CKO-CPC compared to male WT-CPC whereas no differences in the EZH2 expression in female CPC were observed. Clonally expanded CPC can differentiate into endothelial cells or into adipocytes depending on the differentiation conditions. ZFP423 overexpression is sufficient to induce white adipocyte differentiation of clonal CPC. In isolated WT-CPC, PGD_2_ stimulation reduces the expression of EZH2 thereby upregulating ZFP423 expression and promoting white adipocyte differentiation.

Thus, cardiomyocyte STAT3-deficiency leads to age-related and sex-specific cardiac remodeling and failure in part due to sex-specific alterations in PGD_2_ secretion and subsequent epigenetic impairment of the differentiation potential of CPC. Causally involved is the impaired AR signaling in absence of STAT3, which reduces the expression of the PG degrading enzyme HPGD.

## Introduction

Men and women experience quite different cardiovascular disease susceptibility profiles and outcome, a feature that is poorly understood. Further, the effects of biologic sex on health, disease susceptibility and mortality are vastly understudied (1, 2). Recent studies showed that genetics contribute to sex-specific differences in fat tissue and cardiovascular and metabolic diseases (3). Pathophysiologically enhanced cardiac fat content is frequently observed in patients with heart failure, in arrhythmogenic right ventricular dysplasia (ARVD), and after myocardial infarction (4–6). In patients with dilated cardiomyopathy (DCM), increased fat deposits are associated with more severe left ventricular (LV) dilatation and decreased systolic LV function, compared with DCM patients without enhanced cardiac fat (6). Three different types of fat tissue exist of which brown adipose tissue (BAT), mostly present in embryonic and fetal stages, and beige adipose tissue (BET) present postnatally, utilize glucose and lipids to generate heat and are associated with improved cardiometabolic health (7). Brown and beige adipocytes harbor many similar properties, including multilocular lipid droplets, dense mitochondria, and the activation of a thermogenic gene program involving the PR domain containing 16 (PRDM16) protein. PRDM16 is a transcriptional coregulator that controls the development of brown and beige adipocytes and leads to the upregulation of uncoupling protein 1 (UCP1), a hallmark of BAT/BET (8, 9). The third fat type is defined as white adipose tissue (WAT) which stores energy by accumulating fat droplets. WAT is needed as mechanical protection for organs and secretes cytokines and hormones. Extensive WAT formation is associated with an increased cardiovascular risk as it promotes inflammation and alters the immune-endocrine response (10). WAT-specific gene programs typically upregulate the zinc-finger protein 423 gene (*Zfp423*, the murine ortholog of the human *Znf423*). ZFP423 suppresses the PRDM16-mediated thermogenic program and thereby prevents fat cells from burning energy by keeping white fat cells in an energy-storing state (11). A change in the proportions of adipose tissues occurs with aging, leading to a decline of BAT/BET and an increase in WAT, which plays a central role in cardiovascular diseases and type II diabetes (10, 12). Development, maintenance, and activation of the different adipose tissues are guided by genetic factors and epigenetic programs, which regulate the *de novo* differentiation of adipocytes from progenitor cells, as well as white-to-brown adipocyte transdifferentiation (7). However, little is known about mechanisms driving the build-up of different fat types in the heart, especially under pathophysiological conditions and whether there are sex-specific differences in these processes.

The cardiac expression and activation of the signal transducer and activator of transcription factor-3 (STAT3) diminishes with age and is notably reduced in failing hearts from patients with dilatative cardiomyopathy (DCM) or peripartum cardiomyopathy (PPCM) (13–16). Moreover, cardiomyocyte-specific deficiency of STAT3 (CKO) leads to age-related heart failure and more pronounced cardiac damage and failure in response to ischemic injury and infection in male mice (17–19). Nulli-pari CKO females seem protected from age-related heart failure but develop PPCM after breeding (14). Beside direct protective effects on cardiomyocytes, cardiomyocyte STAT3 influences also the cardiac cell-to-cell communication by regulating the expression and secretion of paracrine factors, impacting on endothelial cells, fibroblasts, inflammatory cells and endogenous stem cell antigen (SCA)-1^+^ cardiac-progenitor-cells (CPC) (17, 20–23).

The present study shows that cardiomyocyte-specific STAT3 deficiency leads to an upregulation of cyclooxygenase-2 (COX-2, also known as prostaglandinsynthase-2, PGHS-2) in young male and female CKO mice thereby promoting the production of the prostaglandin D_2_ (PGD_2_) from arachidonic acid. In males, the prostaglandin degrading enzyme hydroxyprostaglandin-dehydrogenase-15 (HPGD) expression is under the control of the androgen receptor (AR) for which STAT3 acts as a co-factor (24). As a consequence, PGD_2_ secretion from male CKO-CM but not from female CKO-CM is increased which subsequently represses the Enhancer of Zeste homolog 2 (EZH2) subunit of the Polycomb repressive complexe 2 (PRC2), a histone methyltransferase associated with transcriptional repression of the white adipocyte differentiation factor ZFP423 (25). In turn, EPO, which is also reduced in CKO hearts, has been shown to enhance endothelial differentiation from CKO-CPC (22) and the present study shows that it suppresses the ZFP423 expression and white adipocyte differentiation from CKO-CPC.

In summary, STAT3-deficiency leads to sex-specific alterations in the cardiomyocyte secretome which provokes an epigenetic shift in CPC from endothelial cells towards white adipocytes differentiation in male but not female CKO hearts. This process contributes to a decline in capillaries and an increase in WAT deposits with cardiac remodeling and heart failure in aging male but not female CKO mice.

## Results

### Male but not female CKO mice develop age-related heart failure with increased intraventricular fat accumulation and enhanced inflammation and fibrosis

We previously showed that male and female mice with a cardiomyocyte-specific STAT3 deficiency (αMHC-Cre^tg/+^; STAT3^flox/flox^, CKO) exhibit normal cardiac function and morphology at young age (3 months) and that male but not female mice develop left ventricular (LV) systolic dysfunction at 6 months of age (S1 Table and S2 Table) (14, 17). Histological analyses (Oil Red O and Perilipin staining) revealed that LV adipocyte content was low with no difference between WT and CKO mice of both sexes at the age of 3-4 months (Figs 1A and 1B, S1A Fig). At 6-months, however, LVs from male but not female CKO mice displayed increased adipocytes content compared to age- and sex-matched controls (Figs 1A and 1B, S1A Fig). Further analyses revealed positive staining for the adipocyte marker Perilipin and the white adipocyte marker Resistin while the brown/beige adipocyte marker UCP1 could not be detected (Fig 1C, S1B Fig). In addition, the triglyceride content was higher in LV tissue from 6-month-old male CKO hearts compared to age-matched male WT hearts (Fig 1D). Aged male CKO hearts displayed reduced capillary density, elevated numbers of inflammatory cells (CD45 positive infiltrates), increased expression of the monocyte/macrophage marker ADGRE1 and increased fibrosis compared with hearts from age-matched male WT mice (Figs 1E–1H).

**Fig 1.**
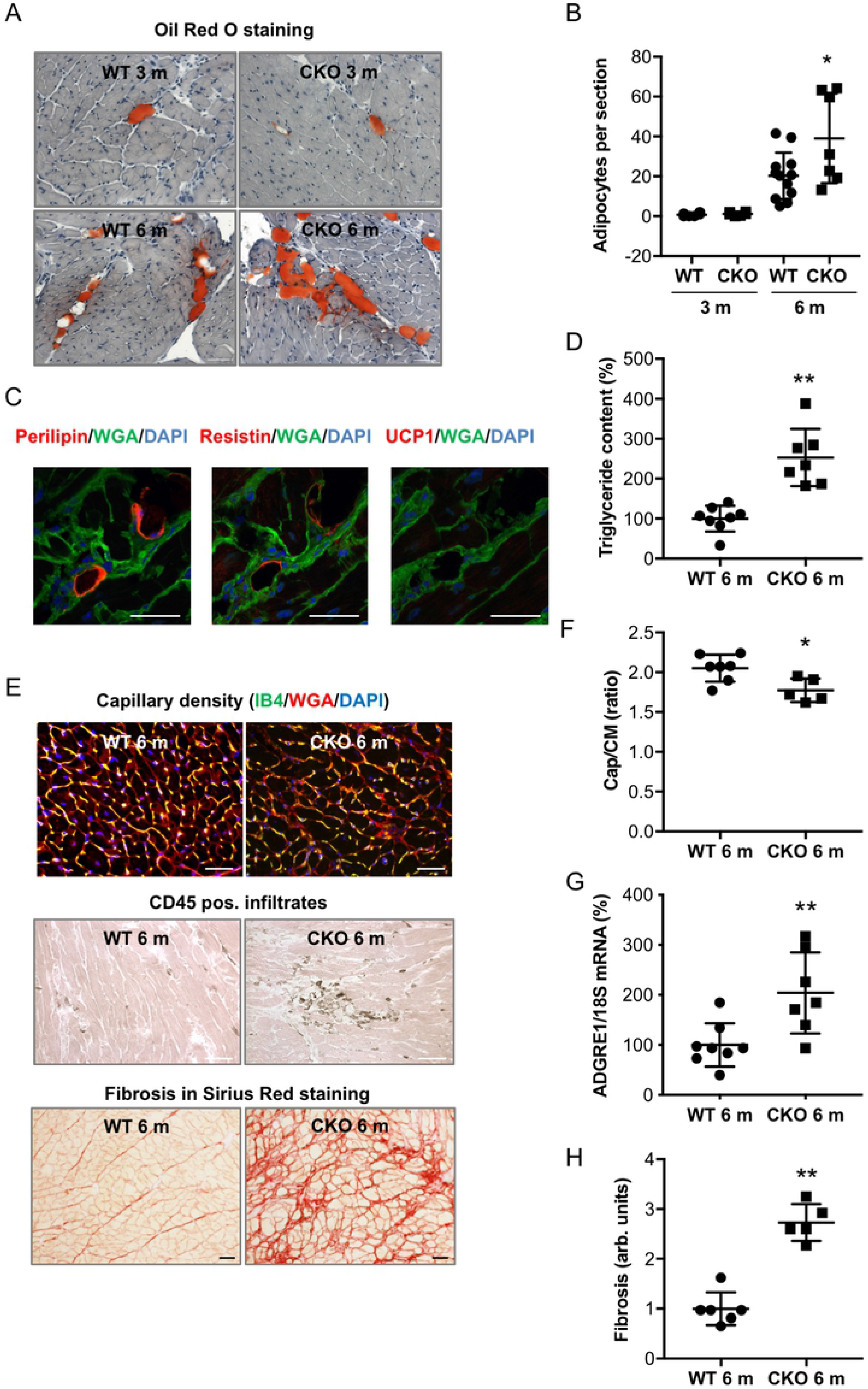
Male CKO mice with age-related heart failure display increased intraventricular fat accumulation with enhanced inflammation and fibrosis. **(A)** Oil Red O staining of adipocytes in LV cryosections counterstained with hematoxylin of 3- or 6-month-old (m) male WT or CKO mice, scale bars: 50 μm. **(B)** Bar graph summarizes the number of adipocytes per section in 3- or 6-month-old male WT (3 m: n=6; 6 m: n=12) and CKO (3 m: n=6; 6 m: n=7) LVs, *P<0.05 vs. WT 6 m, two-way ANOVA with Bonferroni’s multiple comparison test. **(C)** Immunofluorescence staining of Perilipin (red), Resistin (red) or UCP1 (red) counterstained with WGA-FITC (green) and DAPI (blue) in cryosections of heart tissue (male 6 m CKO mice), scale bars: 25 μM. **(D)** Quantification of triglyceride content in LV extracts from 6 m male WT (n=8) and CKO (n=7) mice, mean of WT was set at 100 %. **(E)** Capillary density (upper panel: isolectin B4 (IB4, green)/WGA (red) and nuclear staining DAPI (blue), CD45 positive infiltrates (middle panel: brown, co-stained with eosin) and Sirius Red staining (lower panel) of LV cryosections of 6 m male WT or CKO mice, scale bars: 50 μm. **(F)** Capillary density determined as the ratio of capillaries to cardiomyocytes in transversely sectioned male WT (n=7) and CKO (n=5) LVs. **(G)** Dot plot summarizes ADGRE1 mRNA levels of male WT (n=8) and CKO LVs (n=7), mean of WT was set at 100 %. **(H)** Quantification of fibrosis from WT (n=6) and CKO (n=5) LVs in arbitrary units (arb. units). **(D-H)** All data are mean±SD, *P<0.05, **P<0.01 vs. WT, two-tailed unpaired t-test.

### COX-2 expression is increased in male and female CKO cardiomyocytes but reduced HPGD expression and increased PGD_2_ secretion are only present in male cardiomyocytes

Prostaglandins are metabolites of the arachidonic pathway in which COX-1 and COX-2 and HPGD are rate limiting enzymes. LV tissue from 3- and 6-month-old CKO male mice expressed less HPGD than WT male mice (Fig 2A, S2A Fig) while no such difference was visible between CKO- and WT female mice (S2B and S2C Figs). Moreover, HPGD expression was also lower in male CKO- compared to male WT-CM (Fig 2B) while female CKO- and WT-CM showed no difference (S2D Fig).

**Fig 2.**
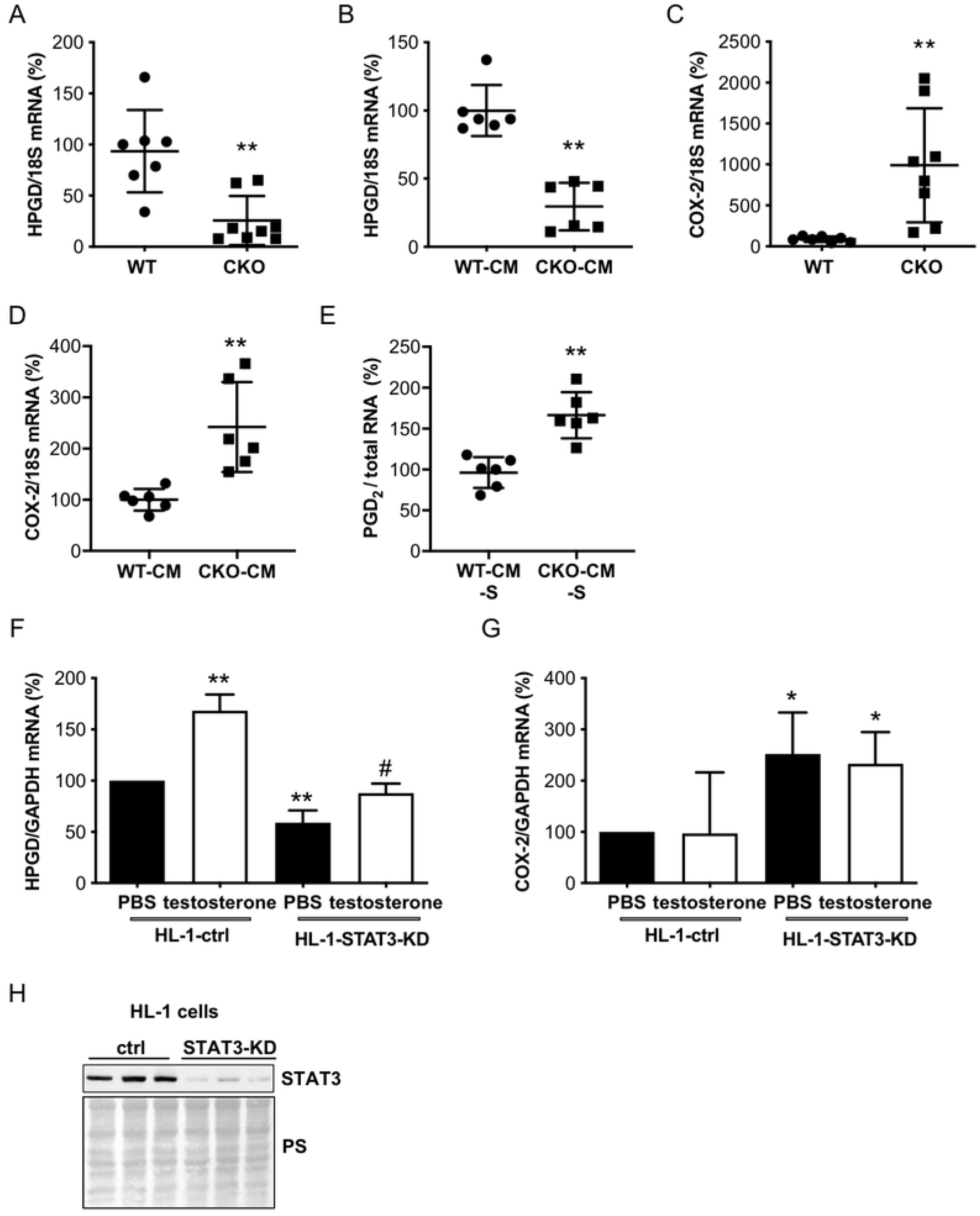
STAT3 deficiency alters COX-2 and HPGD expression in male cardiomyocytes leading to increased prostaglandin D_2_ levels. **(A, C)** Dot plots summarize mRNA levels of **(A)** HPGD and **(C)** COX-2 in LVs of 3-month-old male WT (n=7) and CKO mice (n=8). **(B, D)** Dot plots summarize **(B)** HPGD and **(D)** COX-2 mRNA levels of WT- and CKO-cardiomyocytes (CM) isolated from 3-month old mice (n=6 animals per genotype). **(E)** Measurement of PGD_2_ levels in supernatants of male WT- and CKO-CM assessed by ELISA (n=6 animals per genotype), normalized to total RNA concentrations. **(F, G)** Bar graphs summarize mRNA levels assessed by qRT-PCR of **(F)** HPGD and **(G)** COX-2 in HL-1 cells treated with testosterone (10 nM) for 24 h. **(H)** Representative western blot showing protein expression of STAT3 in HL-1 control (ctrl) and STAT3-KD cells. Ponceau S (PS) served as a loading control. **(A-E)** All data are mean±SD, WT mean was set at 100 %, **P<0.01 vs. WT, two-tailed unpaired t tests. **(F-G)** Data are presented as mean±SD, n=4, mean of HL-1-ctrl PBS was set at 100 %, *P<0.05, **P<0.01 vs. HL-1-ctrl PBS, ^#^P<0.05, ^##^P<0.01 vs. HL-1-STAT3-KD PBS, two-way ANOVA with Bonferroni’s multiple comparison test.

Increased COX-2 expression was observed in LV tissue of 3- and 6-month-old male and female CKO mice and in isolated male and female cardiomyocytes from young (3-month-old) CKO mice compared to respective sex- and age-matched WT controls (Figs 2C and 2D, S2E-H Figs). PGD_2_ levels were increased in the supernatants of male CKO-CM compared to levels in the supernatants of male WT-CM (Fig 2E). Female CKO-CM displayed no differences in secreted PGD_2_ levels compared to female WT-CM (S2I Fig).

### Testosterone-induced expression of HPGD is attenuated in STAT3-knockdown HL-1 cardiomyocytes

It has been reported that HPGD is positively regulated by the androgen receptor (AR) (26) and that the AR is expressed by HL-1 cardiomyocytes (27). We observed that HPGD expression was significantly lower in control HL-1-STAT3-KD-CM compared to HL-1-ctrl-CM (Figs 2F and 2H). Testosterone stimulation markedly induced HPGD in HL-1-ctrl-CM (+68±16%), which was attenuated in HL-1-STAT3-KD-CM (+29±9%) (Fig 2F). In turn, COX-2 was higher in HL-1-STAT3-KD-CM without an additional effect of testosterone treatment compared to HL-1-ctrl-CM (Fig 2G). Treatment with estrogen did not influence HPGD or COX-2 expression of HL-1-ctrl- or HL-1-STAT3-KD-CM (S2J and S2K Figs).

### PGD_2_ promotes white adipocyte differentiation of WT-CPC and of human induced pluripotent stem cells

PGD_2_ upregulated white adipocyte differentiation in WT-CPC as shown by increased Oil Red O positive cells, which was associated with an early reduction of the Enhancer of Zeste homolog 2 (EZH2) subunit of the Polycomb repressive complex 2 (PRC2), a histone methyltransferase associated with transcriptional repression of the white adipocyte differentiation factor ZFP423 (25) and enhanced ZFP423 expression (Figs 3A–3D). PGD_2_ also enhanced expression levels of the adipocyte markers PPARγ, CCAAT enhancer binding protein alpha (CEBPA) and fatty acid binding protein 4 (FABP4) and of the WAT markers ZFP423, lysozyme 2 (LYZ2) and Resistin, while the expression of BAT/BET markers (EBF transcription factor-2, EBF2 and transmembrane protein 26, TMEM26) remained unchanged (Figs 3E–3L) and PR domain containing 16 (PRDM16) and uncoupling protein 1 (UCP1) expression was not detectable. PGD_2_ treatment induced also white adipocyte differentiation in human induced pluripotent stem cells (iPSC) as shown by Oil Red O and Resistin positive cells (S3A Fig). In addition, ZNF423 and CEBPA expression were elevated and EZH2 expression was reduced in PGD_2_ treated human iPSC compared to control iPSC (S3B–S3D Figs).

**Fig 3.**
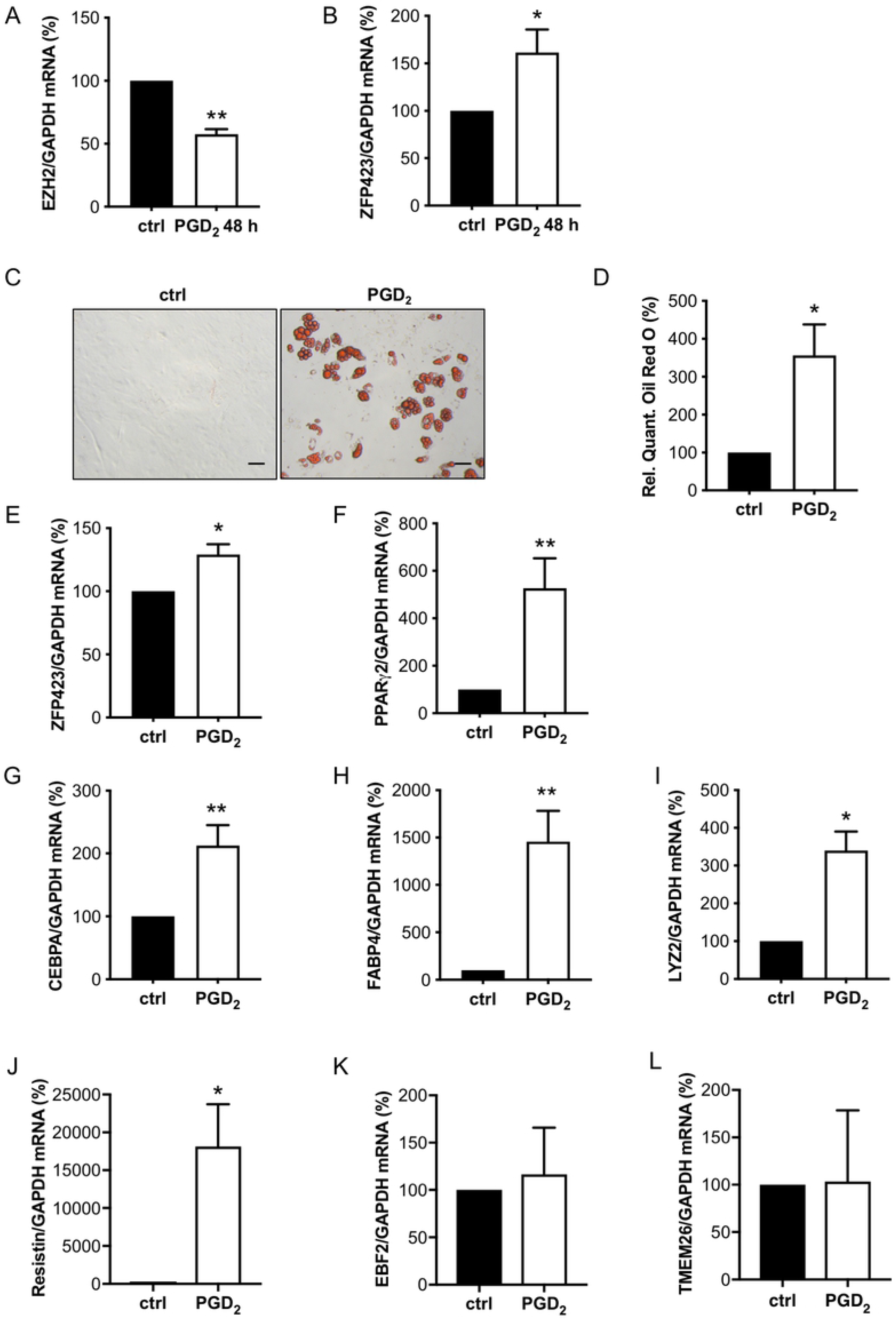
PGD_2_ promotes white adipocyte differentiation of male WT-CPC. **(A, B)** Bar graphs summarize mRNA levels assessed by qRT-PCR of **(A)** EZH2 and **(B)** ZFP423 in WT-CPC after treatment with PGD_2_ for 48 h. **(C)** Representative picture after Oil Red O staining of isolated WT-CPC treated with PGD_2_ (1 μM) for 48 h and cultivated for 12 days, scale bars: 50 μm. **(D)** Relative quantification of Oil Red O measured by absorbance at 492 nm. **(E-J)** Bar graphs summarize mRNA levels assessed by qRT-PCR of **(E)** ZFP423, **(F)** PPARγ2, **(G)** CEBPA and **(H)** FABP4 in WT-CPC incubated with PGD_2_ (1 μM) for 48 h and cultivated for 12 days. **(I-L)** Bar graphs summarize mRNA levels assessed by qRT-PCR of the white adipocyte markers **(I)** LYZ2 and **(J)** Resistin and of the brown/beige adipocyte markers **(K)** EBF2 and **(L)** TMEM26 in WT-CPC incubated with PGD_2_ (1 μM). **(A, B, I-L)** Data are presented as mean±SD (WT-CPC isolated and pooled from 6 animals), mean of control cells was set to 100 %, *P<0.05, **P<0.01 vs. control, one sample t-test. **(E-H)** Data are presented as mean±SD (WT-CPC isolated and pooled from 3 animals), mean of control cells was set to 100 %, *P<0.05, **P<0.01 vs. control, one sample t-test.

### CPC isolated from young male CKO mice display increased adipocyte differentiation potential compared with CPC isolated from young male WT mice

Based on the higher secretion of PGD_2_ from male CKO-CM and the effect of PGD_2_ on CPC from WT mice (WT-CPC), we evaluated the differentiation potential of freshly isolated CPC from young WT and CKO male mice for which we previously showed that STAT3 is exclusively deleted in cardiomyocytes of CKO mice, while its expression is comparable in CPC from CKO (CKO-CPC) and WT mice (22). CPC showed marked expression of platelet-derived growth factor receptor (PDGFR)α indicative for their mesenchymal stem cell character (Fig 4A and 4B). The lack of preadipocyte factor 1 (PREF-1, highly expressed in the preadipocyte cell line 3T3-L1) expression in CPC confirms that isolated CPC were not contaminated with preadipocytes (Figs 4A and 4B). *In vitro* cultivation led to spontaneous differentiation into various cell types including endothelial cells and adipocytes, as shown previously (22) (Figs 4C and 4D). The spontaneous adipocyte differentiation was 2-fold higher in male CKO-CPC compared with male WT-CPC, despite identical cultivation conditions (Figs 4D and 4E), while no elevated adipocyte differentiation was observed in female WT-CPC and CKO-CPC. After differentiation, male CKO-CPC displayed an upregulation of the general adipocyte markers CEBPA and FABP4 (Figs 4F and 4G), and WAT markers ZFP423, LYZ2 and Resistin (Figs 4H–4J). The expression of the BAT/BET markers EBF2 and TMEM26 was similar in CKO- and WT-CPC (Figs 4K and 4L) and the BAT/BET markers PRDM16 and UCP1 could not be detected.

**Fig 4.**
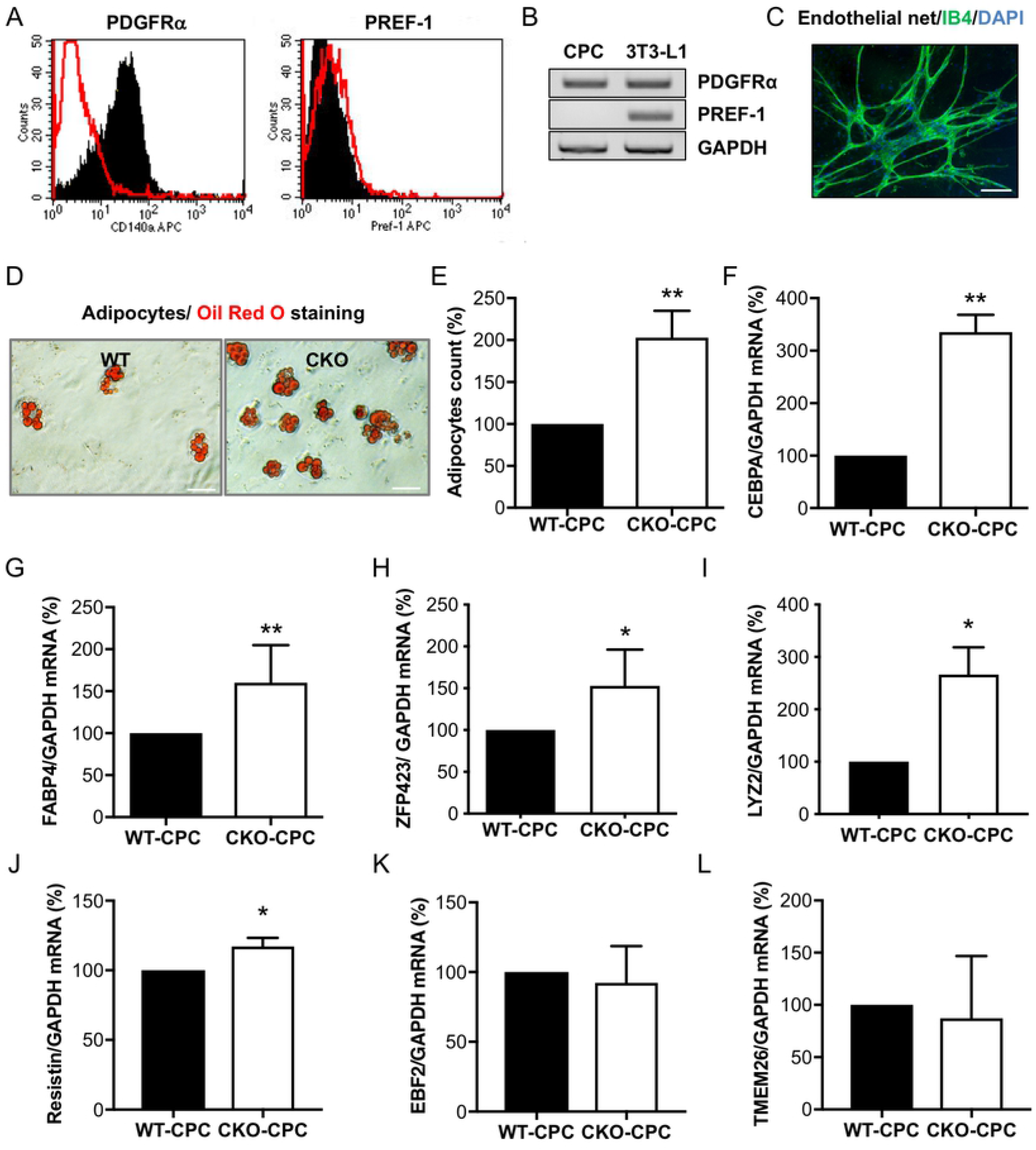
Characterization and adipocyte formation of cultivated CPC isolated from young CKO and WT male mice with normal cardiac function. **(A)** Flow cytometry (PDGFRα or PREF-1: black; IgG control: red) and **(B)** qRT-PCR analysis of PDGFRα and PREF-1 in freshly isolated male WT-CPC, n=3 independent isolations (each isolation consists of 8-12 animals). The preadipocyte cell line 3T3-L1 served as a positive and GAPDH as a loading control. **(C)** IB4 staining of WT-CPC after 4 weeks in culture (IB4, green; DAPI, blue; scale bar: 100 μm). **(D)** Oil Red O staining visualizes spontaneous differentiation of adipocytes in WT- and CKO-CPC after 4 weeks of cultivation, scale bars: 50 μm. **(E)** Bar graph summarizing adipocyte counts of Oil Red O positive cells (n=5 independent isolations; each isolation consists of 10-12 animals per genotype). **(F-H)** QRT-PCR detects mRNA levels of adipocyte markers **(F)** CEBPA, **(G)** FABP4 and **(H)** ZFP423 after 4 weeks of cultivation in WT- and CKO-CPC (n=3, independent cell isolations; each isolation consists of 10-12 animals per genotype). **(I, J)** QRT-PCR detects mRNA levels of WAT markers **(I)** LYZ2 and **(J)** Resistin after *in vitro* cultivation in WT- and CKO-CPC (n=3 isolations per genotype). **(K, L)** QRT-PCR detects mRNA levels of BAT/BET markers **(K)** EBF2 and **(L)** TMEM26 after *in vitro* cultivation in WT- and CKO-CPC (n=3 isolations per genotype). **(E-L)** Bar graphs represent mean±SD, mean of WT was set at 100 %, *P<0.05, **P<0.01 vs. WT, one sample *t*-test.

### The epigenetic signature of freshly isolated CPC isolated from young CKO males differs from CPC isolated from young WT males

Since adipocyte priming of CKO-CPC was maintained during *in vitro* cultivation, we analyzed the epigenetic profiles of CKO-CPC and WT-CPC freshly isolated from 3-month-old male mice with normal cardiac function (S1 Table). A genome-wide exploratory methylation profiling by RRBS, revealed a total of 83 differentially methylated regions (DMRs) overlapping with 81 unique genes with a minimum group methylation difference of 0.1 and p<0.001 (S4A Fig). DMRs with hypermethylated regions were detected in genes such as *Epas1* and *Fgfr2*, both known to promote angiogenesis (28, 29) (S4A Fig, Figs 5A and 5B). In turn, regions in the vicinity of the *Zfp423* 5-UTR (in exon 2, chr8: 87783293-87783352, Wilcoxon p<5.5e-6) were hypomethylated in CKO-CPC compared with WT-CPC (S4A Fig, Fig 5C). *Zfp423* expression is negatively regulated by zinc finger protein 521 (ZFP521) (30), which was slightly hypermethylated in CKO-CPC (Wilcoxon p<0.02, S4A Fig, Fig 5D). QRT-PCR confirmed higher mRNA expression of ZFP423 in CKO-CPC compared with WT-CPC, while no difference was observed for ZFP521 expression (Figs 5E and 5F). *Zfp423* expression can also be regulated by bone morphogenetic protein (BMP)2 and BMP4 (30, 31), but no differences were detected in the methylation pattern of these genes and the expression of BMP2 and BMP-4 in LV heart tissue taken from CKO and WT male mice was similar (Figs 5G and 5H). EZH2 expression was reduced in freshly isolated CKO-CPC compared with WT-CPC from male mice (Fig 5I). In turn, no alteration in EZH2 or ZFP423 mRNA levels were observed in CKO- and WT-CPC isolated from 3-month-old female mice (Figs 5J and 5K).

**Fig 5.**
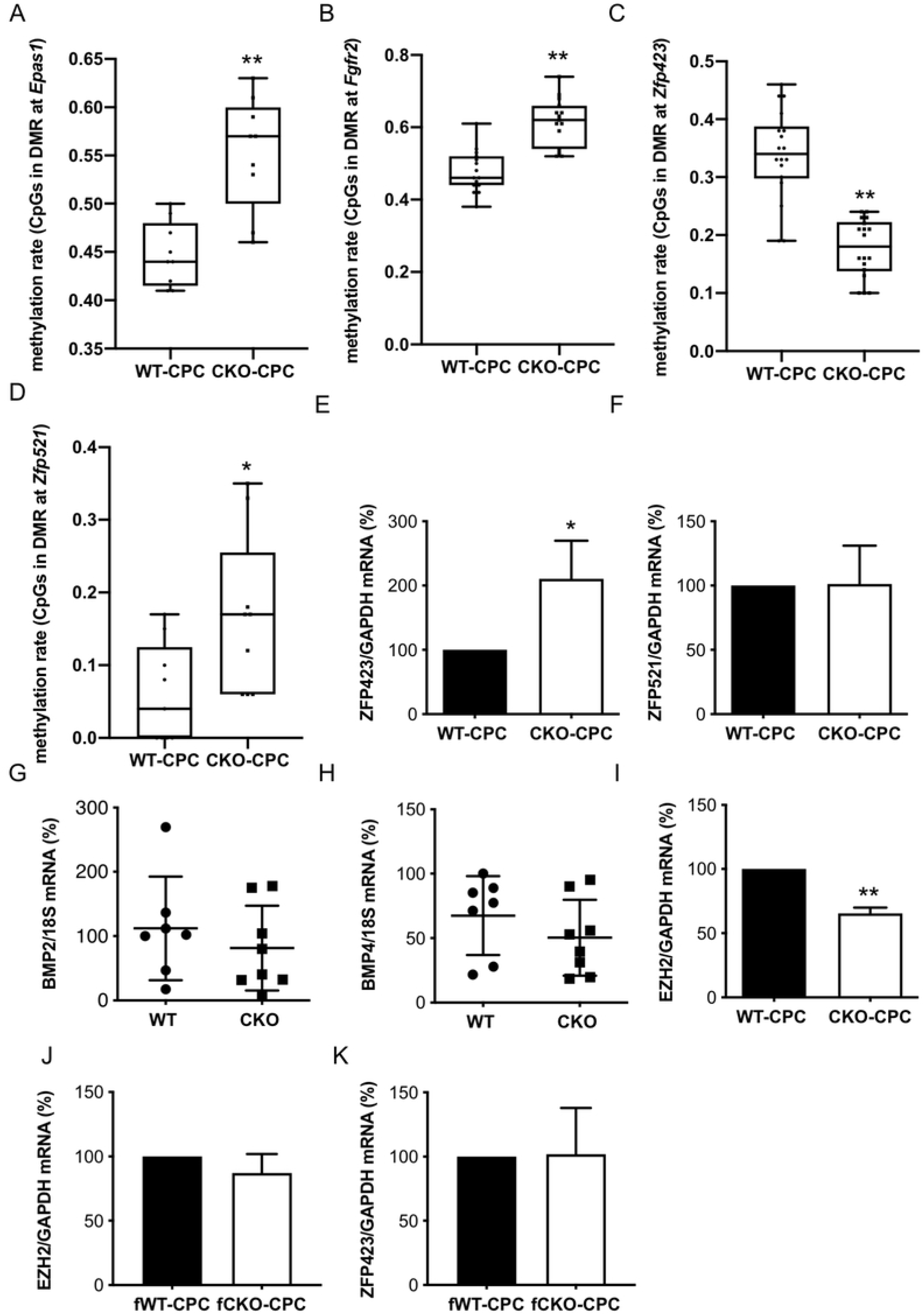
Epigenetic analysis of freshly isolated CKO- and WT-CPC. **(A, B)** Boxplots with single CpG methylation values for **(A)** *Epas1* (chr17:86759066-86759095) and **(B)** *Fgfr2* (chr7:130728553-130728597). **(C)** Boxplot with single CpG methylation values in a region overlapping with the 5’-UTR of exon 2 of *Zfp423* in CKO- and WT-CPC (chr8: 87783293-87783352). Each group contains 3 samples with methylation values for 6 CpGs within the indicated genomic interval, respectively (n=18). **(D)** Boxplot with single CpG methylation of a region approximately 1.5kb downstream of the second exon of *Zfp521* in CKO- and WT-CPC (chr18:13969640-13969667). For each of the 3 samples per group, 3 CpG methylation values in the identified region (n=9) are shown. **(E, F)** Bar graphs summarize mRNA levels detected by qRT-PCR of **E** ZFP423 and **F** ZFP521 in freshly isolated WT- and CKO-CPC (n=5 independent isolations, each isolation consists of 8-12 animals). **(G, H)** Dot plots summarize **(G)** BMP2 and **(H)** BMP4 mRNA levels of WT (n=7) and CKO (n=8) LVs. **(I)** Bar graph summarizes EZH2 mRNA levels detected by qRT-PCR in freshly isolated WT-36 and CKO-CPC from 3-month-old male (n=5 independent isolations, each isolation consists of 8-12 animals). **(J, K)** Bar graphs summarize **(J)** EZH2 and **(K)** ZFP423 mRNA levels detected by qRT-PCR in freshly isolated CPC from 3-month-old female WT and CKO mice (CPC isolated and pooled from 3 WT and 2 CKO mice). **(A-D)** Differences in methylation values were tested by Wilcoxon test, *P<0.05, **P<0.001. **(E-K)** Data are presented as mean±SD, mean of WT was set to 100 %, *P<0.05 vs. control, **P<0.01 vs. control; (**E, F, J-K)** one sample *t*-test and (**G, H)** two-tailed unpaired t test.

### Clonally expanded CPC have the potential to differentiate into endothelial cells and white adipocytes

As freshly isolated CPC are generally a heterogenic pool of cells (22), two clonally expanded CPC cell lines (cCPC) (22, 32) were tested regarding their potential to differentiate into endothelial cells and adipocytes. We observed that the same passage of cCPC cell lines differentiate into endothelial cells on Matrigel (22, 32) or into adipocytes when adipocyte differentiation media and the peroxisome proliferator activated receptor γ (PPARγ) activator indomethacin (33) were added (S5A and S5B Figs). Adipocyte differentiation led to a reduction in SCA-1 and PDGFRα expression and an upregulation of the general adipocyte markers CEBPA and FABP4 (S5C–S5F Figs). Further analyses showed upregulation of the WAT markers LYZ2, and Resistin while the expression of the BAT/BET markers EBF2 and TMEM26 remained unchanged (S5G–S5J Figs) and PRDM16 and UCP1 were undetectable. Retroviral overexpression of ZFP423 induced white adipocyte differentiation in cCPC as shown by increased Oil Red O staining and enhanced expression of PPARγ2, CEBPA, LYZ2 and Resistin (Figs 6A–6G). The expression of the BAT/BET markers EBF2 and TMEM26 remained unchanged (Figs 6H and 6I) and PRDM16 and UCP1 were not detectable.

**Fig 6:**
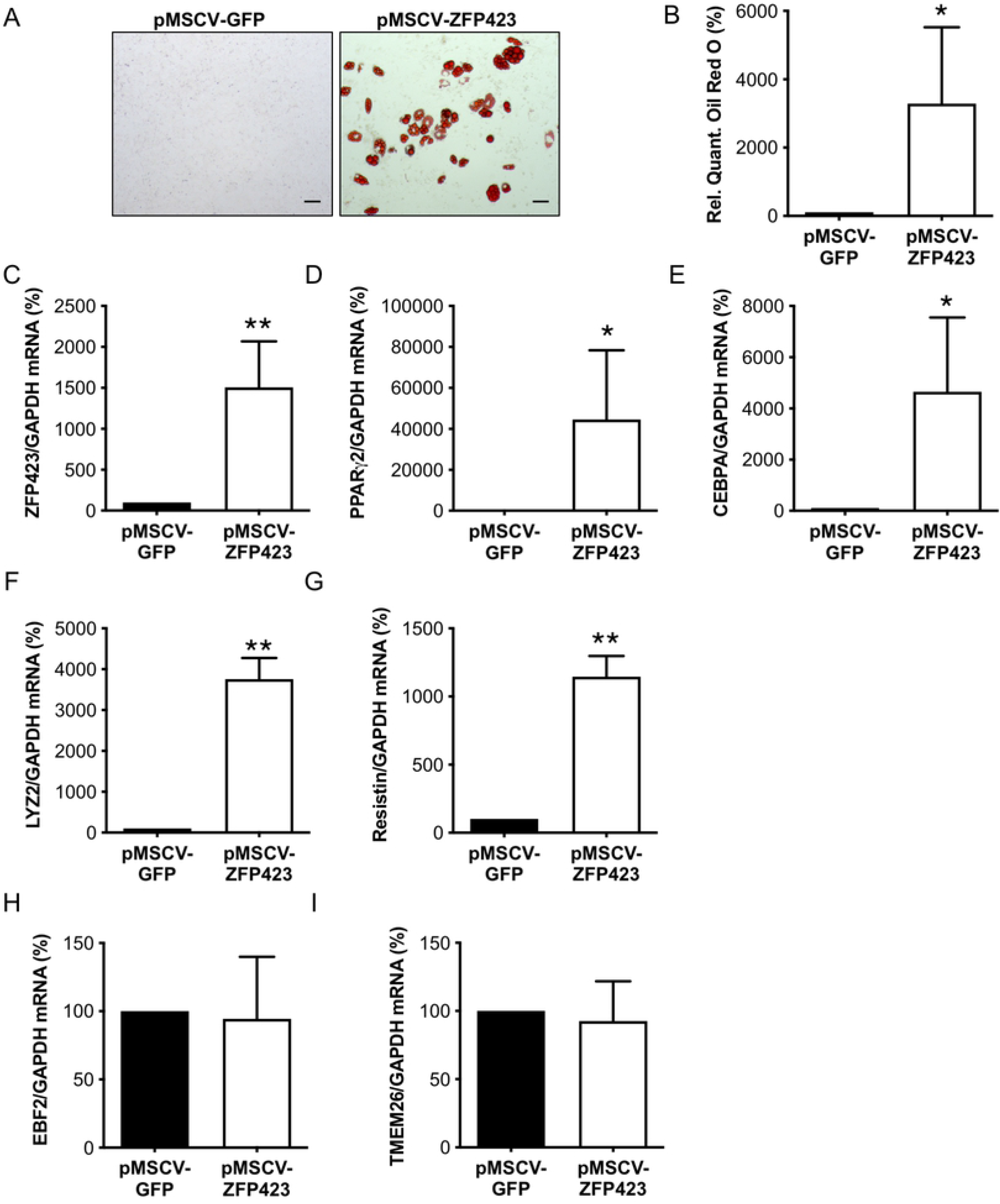
Retroviral overexpression of ZFP423 leads to white adipocyte differentiation of cCPC. **(A)** Oil Red O staining of cCPC with retrovirally-mediated overexpression of ZFP423. Control cells were transduced with the pMSCV-GFP control virusplasmid, scale bars: 50 μm. **(B)** Relative quantification of Oil Red O measured by absorbance at 492 nm. **(C-E)** QRT-PCR detects mRNA levels of **(C)** ZFP423 and of the adipocyte markers **(D)** CEBPA and **(E)** PPARγ. **(F, G)** qRT-PCR visualizes mRNA levels of the white adipocyte markers **(F)** LYZ2 and **(G)** Resistin. **(H, I)** qRT-PCR visualizes mRNA levels of the brown/beige adipocyte markers **(H)** EBF2 and **(I)** TMEM26. **(B-I)** Data are presented as mean±SD, mean of pMSCV-GFP transduced control cells was set to 100 % (n=3 independent cell culture experiments), *P<0.05, **P<0.01 vs. pMSCV-GFP transduced control cells, one sample *t*-test.

### EPO reduces ZFP423 expression and adipocyte differentiation of CKO-CPC

We previously demonstrated that the addition of recombinant murine (rm)EPO to CKO-CPC cultures restored their endothelial differentiation potential (22). Here, we observed that addition of rmEPO to CKO-cultures persistently reduced the ZFP423 expression in CKO-CPC (S6A and S6B Figs), which was associated with attenuated adipocyte differentiation, as shown by reduced Oil Red O positive cells and reduced expression of CEBPA and FABP4 (S6C–S6F Figs). However, addition of rmEPO to isolated CPC for 48 h had no effect on EZH2 expression (S6G Fig).

### LV tissue from male patients with end-stage heart failure due to DCM/ICM display higher cardiac CEBPA expression and lower HPGD expression compared with non-failing LV samples from healthy male organ donors

We previously showed that STAT3 is reduced in hearts from patients with end-stage heart failure (15). Here, we observed that LV tissue from male patients with end-stage heart failure due to DCM or ischemic cardiomyopathy (ICM) (n=8) displayed higher CEBPA (+133±142%, P<0.05) and a non-significant trend to higher COX-2 (+33±91%, n.s.) expression compared with LV samples from healthy male organ donors (n=6). In addition, HPGD mRNA levels were significantly lower in male failing LV samples (−55±12%, P<0.05) compared with LV samples from healthy male organ donors.

## Discussion

We previously reported that cardiomyocyte-specific STAT3-deficiency in male but not female mice leads to age-related heart failure (14, 17). Here, we provide evidence that impaired AR receptor signaling caused by STAT3 deficiency leads to a higher secretion of PGD_2_ by male but not female CKO-CM (Fig 7). PGD_2_ subsequently impairs the vascular regeneration potential of CPC by inducing an epigenetic shift from endothelial to white adipocyte differentiation and thereby leads to a degradation of capillaries and an increase in WAT deposits in hearts from aging male but not female CKO mice (Fig 7). These data shed light on sex-specific age-related remodeling processes that contribute to age-related heart failure in males.

**Fig 7.**
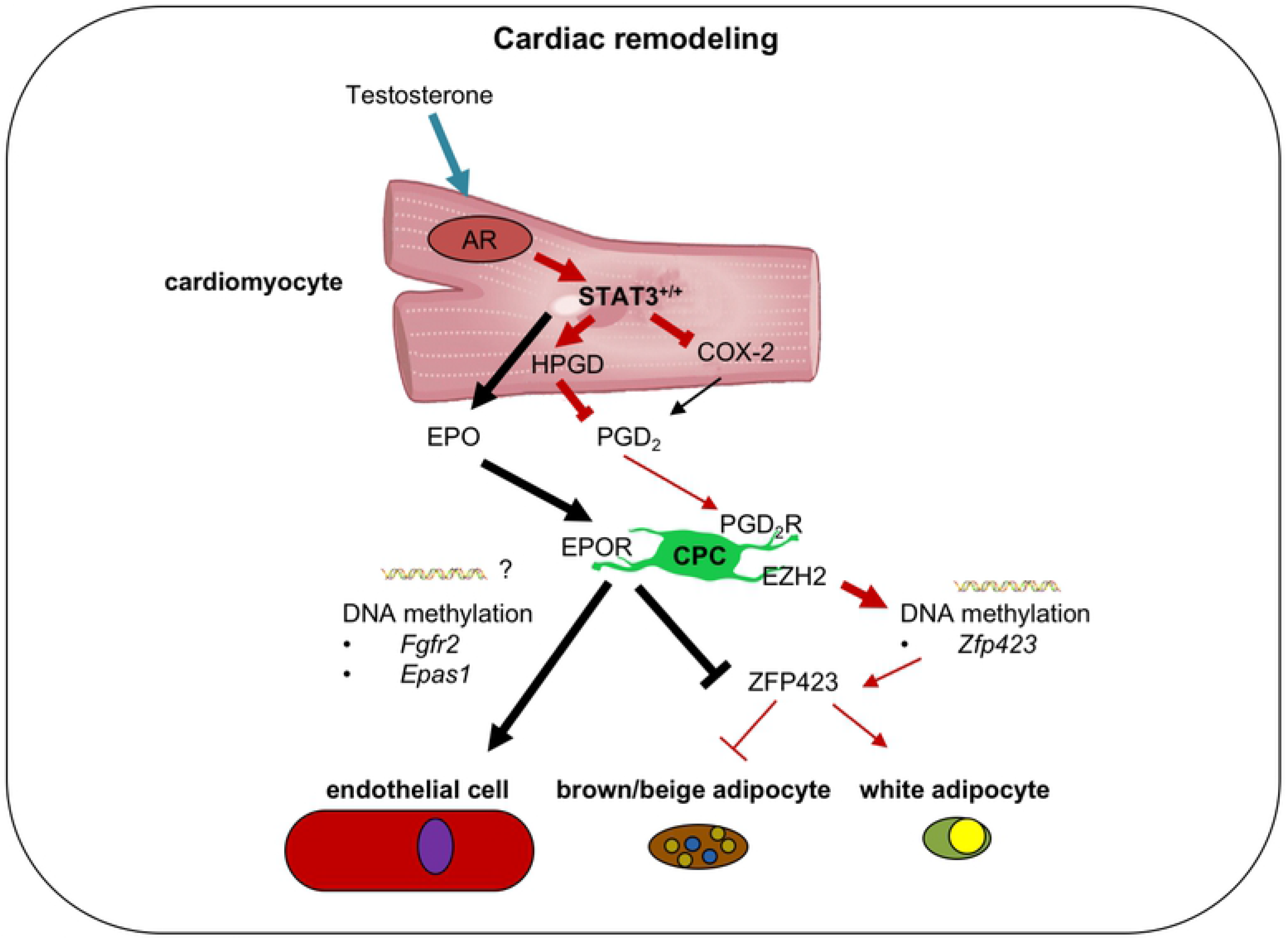
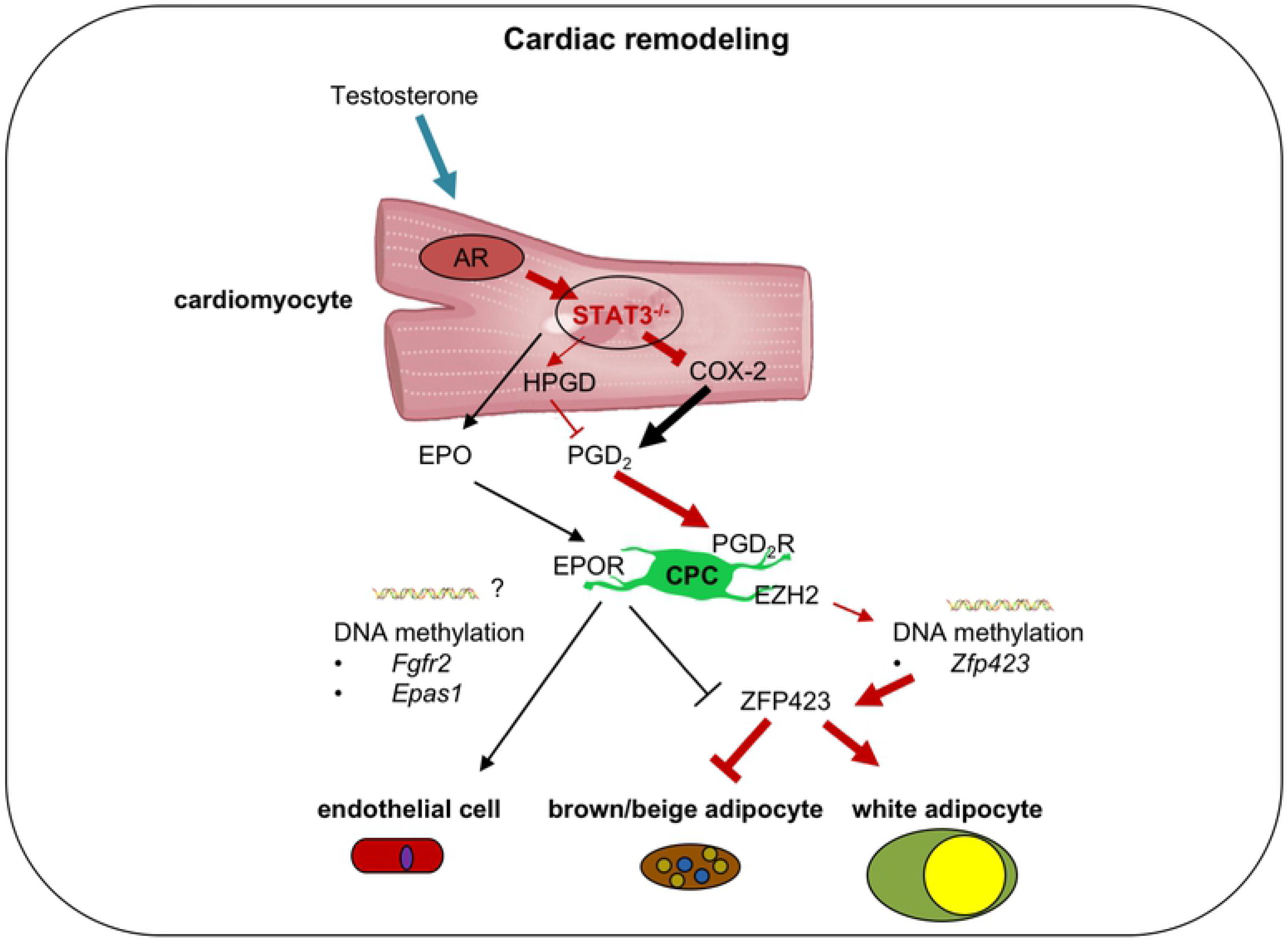
Scheme explaining how cardiomyocyte STAT3-deficiency leads to enhanced white adipocyte differentiation of CPC in male mice contributing to sex-specific cardiac remodeling. The present scheme shows in the upper picture (Fig 7A) the WT male and in the lower picture the CKO male cardiac signaling (Fig 7B). Sex-specific regulated pathways with dark red arrows and sex-unspecific regulated pathways with black arrows. Cardiomyocyte (CM)-specific deficiency of STAT3 (Fig 7B) leads to enhanced expression of COX-2 in male and female mice whereas expression of HPGD is only reduced in male mice likely to result from impaired male-specific hormonal-mediated AR signaling caused by the missing AR co-factor STAT3 in male CKO-CM. This results in increased secretion of PGD_2_ and of its metabolite PGJ_2_ only in male CKO-CM. Subsequently, elevated PGD_2_ levels in the cardiac microenvironment reduced the expression of the EZH2 histone methyltransferase in CPC leading to reduced DNA methylation of its target gene *Zfp423* and with this to enhanced ZFP423 expression. ZFP423 promotes white and suppresses brown/beige adipocyte differentiation of CPC in male CKO hearts. Reduced secretion of EPO from CKO-CM lowers the endothelial differentiation potential of CPC (22) which is associated with hypermethylation of the angiogenic genes *Epas1* and *Fgfr2*. In addition, EPO restores endothelial differentiation and suppresses ZFP423 expression thereby attenuating the endothelial-to-adipocyte shift of CPC.

PGD_2_ is generated by multiple enzymatic steps from arachidonic acid and COX-2 is a rate limiting enzyme in this process (34). COX-2 is elevated in CM from female and male CKO mice, an observation that contrasts with findings showing that STAT3 is a direct transcription factor of COX-2 in ischemic preconditioning, where activation of cardiac STAT3 upregulates COX-2 in the heart (35). However, in our study with young male and female mice canonical STAT3 signaling is not activated suggesting that inactive STAT3 acts either directly or indirectly as a negative regulator of COX-2, a feature that will be explored in future studies. PGs are also regulated by the PG degrading enzyme HPGD (36) and here we observed lower expression of HPGD in male CKO-CM but not in female CKO-CM. In this context it has been shown that the AR can be activated by different ligands including interleukin (IL)-6 and testosterone (37). The AR is expressed in cardiomyocytes from various species including mouse and human (26, 38, 39) and STAT3 acts as a positive co-factor for AR signaling by directly interacting with amino acids 234-558 in the N-terminal domain of the AR (24). The AR is expressed on HL-1 cells and indeed, HPGD expression is lower in HL-1-STAT3-KD-CM compared to ctrl HL-1-CM. Moreover, testosterone induced a marked increase in HPGD in ctrl HL-1-CM, which was blunted in HL-1-STAT3-KD-CM supporting the notion that impaired AR-signaling under low STAT3 condition leads to lower expression of HPGD and with this explains the sex-specific increase in PGD_2_ secretion in male but not female cardiomyocytes from CKO mice (Fig 7).

We demonstrated that PGD_2_ is able to induce white adipocyte differentiation in WT-CPC. In line with this effect of PGD_2_, freshly isolated male CKO-CPC, if kept under the same cultivation conditions, have an increased ability to differentiate into white adipocytes compared to age-matched WT-CPC isolated from males. Furthermore, depending on the differentiation conditions, clonally expanded CPC can differentiate into endothelial cells or into adipocytes suggesting that not differences in the CPC populations but reduced STAT3 expression in cardiomyocytes and subsequent PGD_2_ secretion promotes the epigenetic shift in the differentiation potential of CKO-CPC isolated from male CKO mice. Indeed, freshly isolated CKO-CPC or PGD_2_-treated WT-CPC displayed an increased production of the white adipocyte differentiation factor ZFP423. ZFP423 is a transcription factor controlling preadipocyte determination and maintenance of white adipocyte identity through suppression of the thermogenic gene program (8, 10, 12, 40, 41). ZFP423 overexpression in cCPC was sufficient to induce white adipocyte differentiation confirming the important role of ZFP423 for the adipocyte commitment of CPC. Increased ZFP423 expression in male CKO-CPC was associated with an upregulation of WAT markers (LYZ2, and Resistin) and either unchanged expression (EBF2 and TMEM26) or no expression (PRDM16 and UCP1) of BAT/BET markers. Finally, and in line with the sex-specific difference in PGD_2_ production, CPC isolated from female CKO hearts did neither exhibit increased ZFP423 expression nor enhanced adipocyte differentiation.

BMP2, BMP4 and ZFP521 are known regulators of ZFP423 (30, 31) and ZFP521 suppresses the adipocyte lineage by direct transcriptional repression of ZFP423 (42). In the present study, expression of ZFP521, BMP2 and BMP4 were not altered in male CKO-hearts or in freshly isolated male CKO-CPC compared with male WT hearts or WT-CPC, suggesting that these factors are not responsible for the ZFP423 regulation in male CKO-CPC. In turn, PGD_2_ and its non-enzymatically generated metabolite 15-deoxy-delta(12,14)-prostaglandin-J_2_ (PGJ_2_) were reported to decrease EZH2 mRNA and protein expression (43). EZH2 is a subunit of PRC2 and functions as a histone methyltransferase that is able to control CpG methylation through direct physical contact with DNA methyltransferases (DNMTs) (44). Reduced EZH2 levels in the *Zfp423* promoter are associated with lower histone and DNA methylation and higher expression of the *Zfp423* gene, which predisposes fetal progenitor cells to adipogenic differentiation (45). Indeed, in freshly isolated male CKO-CPC the EZH2 expression was reduced and several DMRs were hypomethylated in the *ZFP423* gene compared to male WT-CPC. Moreover, PGD_2_ stimulation of WT-CPC reduced EZH2 expression and increased ZFP423 expression supporting the notion that PGD_2_ alters the epigenetic signature of CPC towards adipocyte differentiation. This observation extends previous studies reporting that PGD_2_ suppresses lipolysis in adipocytes and is associated with insulin resistance and body weight gain (46). This novel aspect of PGD_2_ is interesting with regard to the controversial roles of PGD_2_ documented for the cardiovascular system. In fact, it has been shown that PGD_2_ has protective features for example in I/R injury (47) but may also have adverse effects, for example by inducing cardiomyocyte death (48) and/or vasoconstriction (49).

Beside the hypomethylated ZFP423 gene, male CKO-CPC displayed hypermethylated DMRs in genes promoting angiogenesis (e.g., *Epas1* and *Fgfr2* (28, 29)) consistent with their previously shown lower endothelial differentiation potential (22). Previous data showed that EPO secretion is reduced in the cardiac microenvironment from male CKO hearts and exogenous EPO restored the endothelial differentiation capacity in male CKO-CPC (22). The present study found that EPO reduced ZFP423 expression and adipocyte formation from CKO-CPC, suggesting that lower EPO levels in CKO hearts may further promote the shift from endothelial to adipocyte differentiation in CKO-CPC. So far, there is no evidence for a sex-specific regulation of EPO since EPO blood levels in patients undergoing androgen deprivation therapy remained unchanged (50).

Previous data reported reduced STAT3 in failing human hearts (15) and the present study reveals significantly lower HPGD expression in LV tissue from male patients with terminal heart failure. Furthermore, PGD_2_ induced ZFP423 expression and adipocyte differentiation in human iPSC. Therefore, a similar mechanism as described for male mice may also exists in humans and may explain sex-specific development of heart failure.

In conclusion, age-related reduction of cardiomyocyte STAT3 leads to impaired AR receptor signaling and subsequent reduced expression of the PG degrading enzyme HPGD (Fig 7). This results in an increased secretion of PGD_2_ from cardiomyocytes in male but not female hearts (Fig 7). Subsequently, PGD_2_ induces an epigenetic shift in CPC already prior to heart failure which hampers vascular regeneration and increases WAT deposits and thereby promotes adverse remodeling and heart failure (Fig 7).

## Material and Methods

Unless otherwise stated, chemicals and reagents were all purchased from Sigma-Aldrich.

### Animal experiments

Mice with a cardiomyocyte-restricted knockout of STAT3 (CKO: αMHC-Cre^tg/+^; STAT3^flox/flox^) and wildtype mice (WT: STAT3^flox/flox^) were generated as previously described (17). Echocardiography was performed on 3- and 6-month-old, lightly sedated mice (isoflurane inhalation 0.5%) using a Vevo 770 (VisualSonics) as described (14).

All animal studies were conducted in accordance with the German animal protection law and with European Communities Council Directive 86/609/EEC and 2010/63/EU for the protection of animals used for experimental purposes. All experiments were approved by the Local Institutional Animal Care and Research Advisory Committee and permitted by the relevant local authority for animal protection.

### Histology and immunostaining

For cardiac morphological analyses, hearts were embedded in Tissue Tek OCT and frozen at −80 °C. Interstitial collagen was analyzed in picro-sirius red F3BA-stained sections (17, 51). Inflammation was determined in LV cryosections with an antibody recognizing CD45 (BD Pharmingen 550539). Capillary density was determined in transversely sectioned LVs by isolectin B4 (Vector), counterstained with wheat germ agglutinin (WGA, Vector) and DAPI for nuclear stain (14, 51). For immunostainings using primary antibody recognizing Perilipin (#9349, Cell Signaling), Resistin (ab119501, abcam) or UCP1 (ab10983, abcam) cryosections were fixed in acetone, were washed 3 times with PBS and blocked with 10 % donkey serum and 0.3 % Triton in PBS for 1 h at room temperature. Cryosections were stained with Perilipin antibody (1:100), Resistin antibody (1:50) or UCP1 antibody (1:100) overnight at 4°C. Cryosections were washed 3 times and incubation with the secondary antibody Cy3-anti-rabbit (1:250, Jackson ImmunoResearch) and counterstaining with WGA was performed for 2 h at room temperature. Nuclei were stained with DAPI Hoechst 33342 (Sigma-Aldrich). Images were acquired with AxioVert200M microscope, Axiovison software 4.8, Axio Observer 7 and Zen 2.6 pro software (Carl Zeiss) and with SP8 Leica Inverted confocal microscope.

### Isolation, characterization and culture of SCA-1^+^ cardiac progenitor cells

Isolation of SCA-1^+^ cells from hearts of 3-month-old mice was performed as described previously (22), and is described in detail in the supplemental methods.

### DNA Isolation for reduced representation bisulfite sequencing (RRBS)

The DNA of freshly isolated CKO- and WT-CPC of 3-month-old male mice was prepared using the DNeasy Blood and Tissue Kit according to the manufacturer’s protocol (Qiagen).

### Reduced Representation Bisulfite Sequencing (RRBS)

RRBS libraries for a total of n=6 samples (3 sample pools for CKO mice, 3 sample pools for WT mice, 10-12 mice per pool) were prepared using the Ovation RRBS Methyl-Seq System 1-16 with TrueMethyl oxBS (NuGen), skipping the oxidation step but otherwise undertaken according to manufacturer’s instructions. The average conversion rate across all samples was estimated to be 97 %. Sequencing was performed on a NextSeq500 instrument at the Sequencing Core Facility of the Fritz-Lipmann-Institute Jena. The input DNA and concentration used for library preparation are both given in S3 Table.

### Data analysis of RRBS

#### Preprocessing

Sequencing yielded between 139M and 171M sequences (Table 1). After clipping using *cutadapt* (52) (version 1.18; parameters --quality-cutoff 20 --overlap 5 --minimum-length 25 -u 7 -a AGATCGGAAGAGC), between 138M and 170M remained (Table 1). Before and after clipping quality was inspected using *fastqc* (v0.11.5).

**Table 1.** Read numbers and mapping statistics.

| sample | raw reads | clipped reads | mapping |
| --- | --- | --- | --- |
| Set1 CKO | 171274320 | 170175455 | 152837610 (89.81%) |
| Set1 WT | 147258040 | 146432432 | 131141572 (89.56%) |
| Set2 CKO | 143418782 | 142616983 | 127924806 (89.7%) |
| Set2 WT | 139188392 | 138358371 | 123566916 (89.31%) |
| Set3 CKO | 158225479 | 157255130 | 141428776 (89.94%) |
| Set3 WT | 157426725 | 156389740 | 140337521 (89.74%) |

#### Read alignment, conversion rates

Using the Bisulfite Analysis Toolkit (BAT; v.0.1) (53–55), reads were aligned with *segemehl* (v0.2.0) (53) to the Mus musculus reference genome GRChm38.p6 using standard parameters of the module ‘BAT_calling’ in mode ‘-F 1’. For the extraction of conversion rates, we used the module ‘BAT_calling’ with *haarz* (v0.1.7) (53, 55) and *samtools* (v1.6). The filtering of resulting vcf files containing the conversion rates was done using the module ‘BAT_filter_vcf’. On average, mapping rates were ~89 % (S3 Table).

#### Calling of differentially methylated regions

The methylation data obtained from the previous step was summarized using ‘BAT_summarize’ after the adaptation of the source code to allow the processing of mice data (BAT_summarize_mouse). Specifically, the module was called by assigning all 3 WT samples to the control group and all 3 CKO samples to the case group (parameters --in1 Set1_WT_CPC.bedgraph, Set2_WT_CPC.bedgraph, Set3_WT_CPC.bedgraph --in2 Set1_CKO_CPC.bedgraph, Set2_CKO_CPC.bedgraph, Set3_CKO_CPC.bedgraph --groups control, case --h1 WT1, WT2, WT3 --h2 CKO1, CKO2, CKO3). The module was run with *circos* (56) (v0.69-6) and bedGraphToBigWig (v4). The calling of differentially methylated regions was carried out with ‘BAT_DMRcalling’ using *metilene* (v0.2-8; parameters -a control -b case -z “-m 3 -M 1000 -d 0.01 -v 0.0” -p 1 -d 0.01 -c 3) (57). In order to increase the sensitivity in the given setup, the minimum amount of CpGs per DMR was reduced to 3, and the distance between CpGs within one DMR was increased to 1kb. Subsequently, the raw *metilene* output was annotated with the extended ENSEMBL annotation for GRCm38.p6 (v.92) using *bedtools intersect* (v2.22.1) (58). For the extension of the gene annotation, all annotated genes were extended at their 5’-end by 5kb to include the promoter region.

A total of 5 DMRs with an FDR-adjusted p-value were reported. Due to the high homogeneity of the samples and the relatively small size, we decided to analyze two sets of regions filtered for two different unadjusted p-value criteria. The first set with a minimum group methylation difference 0.1 and p<0.001 yielded a total of 83 DMRs overlapping with 81 unique genes. To better inspect the CpG methylation within predicted DMRs (n=83), we calculated the median methylation for the respective DMRs for each sample (sFig. 1c) and visualized the data using R’s pheatmap function (R version 3.4.1, pheatmap version 1.0.12) with standard parameters for clustering after centering and scaling in the row direction.

Sequencing data of the epigenetic analyses are available under accession number PRJNA602737 in the Sequence Read Archive. Due to the sample size and the exploratory nature of this exercise, we omitted the correction for multiple testing. Pooling the CpG methylation values for all 3 samples in the respective groups was used to confirm or reject methylation differences (Wilcoxon) in CKO-CPC compared with WT-CPC.

### SCA-1^+^ cell cloning

SCA-1^+^ cell cloning was described previously (22, 32). The protocol for clonal expansion of CPC is given in the supplemental methods.

### Retroviral-vector mediated expression of ZFP423

For retroviral-vector mediated expression of ZFP423 the pMSCVFLAG-ZFP423 was used (40). pMSCVFLAG-ZFP423 was a gift from Bruce Spiegelman (Addgene plasmid # 24764; RRID:Addgene_24764). The plasmid pMSCV-GFP was used for transduction of control cells. For generation of lentiviral supernatants, the ecotropic Phoenix Packaging cell line (kindly provided by G. Nolan, Stanford University, Stanford, CA, USA) was used (59). Briefly, 5×10^6^ Phoenix packaging cells were seeded on 10 cm plates in DMEM supplemented with 10 % FBS. On the following day, retroviral constructs (5 μg), together with constructs encoding pGagpol (M57) (10 μg) and pEnv K73 (2 μg), were transiently transfected into Phoenix cells by the calcium phosphate transfection method. Six h after transfection, the media were replaced by DMEM supplemented with 10 % FBS and the cells were cultivated overnight. The media were removed and DMEM/F12 supplemented with 10 % FBS was added and further incubated for 24 h. Supernatants were harvested and filtered through a 40 μm nylon filter and were either directly used for the transduction of CPC or stored at −80 °C. Viral supernatants were added to CPC clones for 6 h in the presence of Polybrene (4 μg/ml). Transduction efficiency based on GFP-expression was controlled in all experiments.

### Adipocyte differentiation of cCPC and isolated WT-CPC

For adipocyte differentiation, cCPC or isolated WT-CPC were seeded on 0.2 % gelatin in DMEM/F12 with 10 % FCS supplemented with ITS (10 μg/ml insulin, 5 μg/ml transferrin, 5 ng/ml sodium selenite), rhFGF-basic (10 ng/ml) and EGF (20 ng/ml). The medium was replaced every 2-3 days. Two days after confluence, adipocyte differentiation was induced in DMEM/F12 supplemented with 10 % fetal bovine serum, 1 μM dexamethasone (G-Biosciences), 0.5 mM methylisobutylxanthine, 1 μg/ml insulin and 1 % penicillin-streptomycin (Thermo Fisher Scientific). Indomethacin (100 μg/ml) or prostaglandin D_2_ (1 μM) were added to the adipocyte differentiation medium. After 2 days, the adipocyte differentiation medium was removed and cells were maintained in DMEM/F12 supplemented with 10 % fetal bovine serum and 1 μg/ml insulin. The cells were harvested in TRIzol (Thermo Fisher Scientific) for RNA isolation or were fixed with 4 % paraformaldehyde for Oil Red O staining.

## Information on stimulation of HL-1 cells are provided in the supplemental methods

### Adipocyte differentiation of human iPSC

The protocols for the generation and for the adipocyte differentiation of human iPSC are given in the supplemental methods.

### Endothelial Differentiation of cCPC on matrigel

The plates were coated with 50 μl/cm^2^ Matrigel Basement Membrane Matrix (Corning) at 37 °C for 30 min. Clonal CPC (50000 cells/cm^2^) were cultivated in EGM-2 complete medium (Lonza) overnight. Cells were fixed using 4 % paraformaldehyde.

### Isolation of murine adult primary cardiomyocytes and cell culture

Murine adult primary cardiomyocytes were isolated and cultivated as described previously (60) from 3-month-old WT and CKO mice. Supernatants of adult primary cardiomyocytes were collected after 48 h, centrifuged at 300×g for 10 min to deplete cell fragments and stored at −80 °C. Cells were harvested in TRIzol (Thermo Fisher Scientific) for RNA isolation.

### Isolation of RNA and qRT-PCR

Total RNA was isolated with TRIzol (Thermo Fisher Scientific) and cDNA synthesis was performed as described previously (22). Real-time PCR with SYBR green dye method (Brilliant SYBR Green Mastermix-Kit, Thermo Fisher Scientific) was performed with the AriaMx Real-Time PCR System (Agilent Technologies) as described (22). Expression of mRNA levels were normalized using the 2-ΔΔCT method relative to GAPDH or 18S. A list of qRT-PCR primers used in this study is provided in the supplemental methods (sTab. 4, 5).

### Immunoblotting

Immunoblots were performed according to standard procedures using SDS-PAGE (17) and the STAT3-antibody (#9139, Cell Signaling),

### Flow Cytometry

5*10^5^ freshly isolated CPC were stained with PREF-1 antibody (AF8277, R&D Systems) and PDGFRα antibody (17-1401-81, eBioscience) for 15 min at room temperature. Flow cytometry was performed using the FACSCalibur (BD Biosciences).

### Immunocytochemistry

Immunostainings using isolectin B4 (Vector Laboratories) were performed according to standard procedures (22). For immunostainings using primary antibody recognizing Perilipin (#9349, Cell Signaling) or Resistin (ab119501, abcam) cells were washed 3 times with PBS and blocked with 10 % donkey serum and 0.3 % Triton in PBS for 1 h at room temperature. Cells were stained with Perilipin antibody (1:100) or Resistin antibody (1:50) overnight at 4°C. Cells were washed 3 times and incubation with the secondary antibody Cy3-anti-rabbit (1:250, Jackson ImmunoResearch) was performed for 2 h at room temperature. Nuclei were stained with DAPI Hoechst 33342 (Sigma-Aldrich). Images were acquired with AxioVert200M microscope and Axiovison software 4.8, Axio Observer 7 and Zen 2.6 pro software (Carl Zeiss).

### Oil Red O staining

Oil Red O staining was performed as previously described (22). A detailed description is provided in the supplemental methods.

### Adipogenesis Detection Assay

Triglyceride accumulation in LV heart tissue was quantified by an adipogenesis detection assay (Abcam ab102513) according to the manufacturer’s protocol.

### PGD_2_ detection in supernatants of adult murine cardiomyocytes

PGD_2_ levels in the supernatants of primary adult murine cardiomyocytes were measured using the prostaglandin D_2_ ELISA kit (Cayman Chemicals No. 512031) according to the manufacturer’s protocols. PG levels in supernatants of primary adult murine cardiomyocytes were normalized to the total RNA content of the cells.

### Patient Data

LV tissues were taken from patients undergoing heart transplantation due to end-stage heart failure caused by dilated (DCM) or ischemic (ICM) cardiomyopathy (DCM/ICM, n=8). LV tissue from donor hearts not suited for transplantation served as controls (NF, n=6). The study was approved by the MHH local ethic committee (Nr. 1833-2013).

### Statistical Analyses

Statistical analysis was performed using GraphPad Prism version 5.0a and 8.1.2 for Mac OS X (GraphPad Software, San Diego, CA, USA). Normal distribution was tested using the D’Agostino normality test. Continuous data were expressed as mean ± SD. Comparison between two groups was performed using one sample *t*-test or unpaired two-tailed t-test. When comparing more than two groups, we used Bonferroni’s multiple comparison test after one/two-way ANOVA testing. A two-tailed *P* value of <0.05 was considered statistically significant.

## Acknowledgements

We thank Martina Kasten, Silvia Gutzke, Birgit Brandt, Iris Dallmann and Tim Kohrn for excellent technical assistance. We thank the research core unit for laser microscopy at the Hannover Medical School and the Core Facility of the Fritz-Lipmann-Institute Jena.

## Data availability

The array data of the epigenetic analyses are available under accession number PRJNA602737 in the Sequence Read Archive (SRA).

## Abbreviations

ANOVA: analysis of variance
AR: androgen receptor
ARVC: arrhythmogenic right ventricular dysplasia
BAT: brown adipose tissue
BMP2: bone morphogenetic protein 2
BMP4: bone morphogenetic protein 4
bpm: beats per minute
BW: body weight
CD: cluster of differentiation
cDNA: complementary deoxyribonucleic acid
CEBPA: CCAAT/enhancer-binding protein alpha
COX: cyclooxygenase
CPC: Sca1^+^-cardiac progenitor cell
DCM: dilated cardiomyopathy
DMEM: Dulbecco’s Modified Eagle Medium
DMR: differentially methylated region
EBF2: early B cell factor 2
EGF: epidermal growth factor
EGM-2: endothelial cell growth medium-2
EPAS1: endothelial PAS domain protein 1
EPO: erythropoietin
EPOR: erythropoietin receptor
EZH2: enhancer of zeste homolog 2
FABP4: fatty acid binding protein 4
FCS: fetal calf serum
FGF: fibroblast growth factor
FGFR2: fibroblast growth factor receptor 2
FITC: fluorescein isothiocyanate
FS: fractional shortening
GLUT4: glucose transporter 4
HBSS: Hank’s balanced salt solution
HPGD: hydroxyprostaglandin-dehydrogenase-15
HW: heart weight
IB4: isolectin B4
ICM: ischemic cardiomyopathy
i.p.: intraperitoneal
ITS: insulin-transferrin, sodium selenite
LV: left ventricle
LVEF: left ventricular ejection fraction
m: month
MACS: magnetic-activated cell sorting
MHC: myosin heavy chain
mRNA: messenger ribonucleic acid
NaCl: sodium chloride
NF: non-failing
PCR2: Polycomb repressive complexe 2
PDGFRα: platelet-derived growth factor receptor alpha
PG: prostaglandin
PPARγ2: peroxisome proliferator-activated receptor gamma isoform 2
PPCM: peripartum cardiomyopathy
PRDM16: PR-domain containing 16
PREF-1: preadipocyte factor 1
qRT-PCR: quantitative real-time polymerase chain reaction
rh: recombinant human
rmEPO: recombinant mouse erythropoietin
RRBS: reduced representation bisulfite sequencing
Sca1: stem cell antigen-1
SD: standard deviation
SDS-PAGE: sodium dodecyl sulfate polyacrylamide gel electrophoresis
STAT3: signal transducer and activator of transcription 3
STAT3-KD: STAT3 knockdown
UCP-1: uncoupling protein-1
UTR: untranslated region
WAT: white adipose tissue
WGA: wheat germ agglutinin
WT: wildtype
Zfp423: zinc-finger protein 423
Zfp521: zinc-finger protein 521
ZNF: zinc nuclear factor

## Supporting information

**S1 Fig. Adipocyte formation in heart tissue of 3- and 6-month-old CKO mice and methylation analysis of freshly isolated CKO- and WT-CPC.**

**(A)** Perilipin staining in LV cryosections of 3- and 6-month-old (m) WT or CKO female (f) mice, Perilipin (red), WGA (green) and DAPI (blue): scale bars: 50 μm**. (B)** Immunofluorescence staining of Perilipin (red), Resistin (red) or UCP1 (red) counterstained with WGA-FITC (green) and DAPI (blue) in cryosections of heart tissue (male 6 m CKO mice), scale: 25 μM.

**S2 Fig. STAT3 deficiency alters COX-2 and HPGD expression in LVs of aged male CKO mice, whereas in females only COX-2 expression is altered resulting in no differences in secreted PGD_2_ levels from isolated adult cardiomyocytes.**

**(A, E)** Dot plots summarize mRNA levels of **(A)** HPGD and **(E)** COX-2 in LVs of 6-month-old male WT (n=8) and CKO mice (n=7). **(B, F)** Dot plots summarize mRNA levels of **(B)** HPGD and **(F)** COX-2 in LVs of 3-month-old female WT (n=7) and CKO mice (n=4). **(C, G)** Dot plots summarize mRNA levels of **(C)** HPGD and **(G)** COX-2 in LVs of 6-month-old female WT (n=6) and CKO mice (n=6). **(D, H)** Dot plots summarize **(D)** HPGD and **(H)** COX-2 mRNA levels of isolated adult female WT- and CKO-cardiomyocytes (CM) (CM isolated and pooled from 3 WT and 2 CKO mice). **(I)** Measurement of PGD_2_ levels in supernatants of isolated adult female WT- and CKO-CM assessed by ELISA (CM isolated and pooled from 3 WT and 2 CKO mice), normalized to total RNA concentrations. **(J, K)** Bar graphs summarize mRNA levels assessed by qRT-PCR of **(J)** HPGD and **(K)** COX-2 in HL-1 cells treated with estrogen (10 nM) for 24 h. **(A-I)** All data are mean±SD, WT mean was set at 100 %, *P<0.05, **P<0.01 vs. WT, two-tailed unpaired t tests. **(J, K)** Data are presented as mean±SD, n=4, mean of HL-1-ctrl PBS was set at 100 %, *P<0.05, **P<0.01 vs. HL-1-ctrl PBS, two-way ANOVA with Bonferroni’s multiple comparison test.

**S3 Fig. PGD_2_ treatment leads to white adipocyte differentiation of human iPSC.**

**(A)** Oil Red O (red) and Resistin (red; DAPI (blue)) staining of PGD_2_ treated human iPSC. **(B-D)** Bar graphs summarize mRNA levels assessed by qRT-PCR of **(B)** EZH2**, (C)** ZNF423 and **(D)** CEBPA of PGD_2_ treated human iPSC (two iPSC clones were tested, n=3 independent cell culture experiments for each clone). Bar graphs represent mean±SD, mean of control cells was set at 100 %, *P<0.05, **P<0.01 vs. ctrl, one sample *t*-test.

**S4 Fig. Methylation analysis of freshly isolated CKO- and WT-CPC.**

**(A)** Heatmap of median methylation values in predicted DMRs (n=83) overlapping with 81 unique genes.

**S5 Fig. Adipocyte differentiation and endothelial cell formation of cCPC expanded from single cells.**

**(A, B)** Differentiation of cCPC expanded from single cells **(A)** on Matrigel (left panel: phase contrast, right panel: Isolectin B4 staining (green); scale bars indicate 100 μm) and **(B)** after adipogenic induction (upper left panel: phase contrast; upper right panel: Perilipin staining (red), nuclear staining, DAPI (blue); lower left panel: Oil Red O staining (red); and lower right panel: Perilipin staining (green), nuclear staining, DAPI (blue), Oil Red O staining (red); scale bars indicate 50 μm). **(C-J)** mRNA levels of progenitor cell markers (**(C)** SCA1 and **(D)** PDGFRα), general adipocyte markers (**(E)** CEBPA, **(F)** FABP4), WAT markers (**(G)** LYZ2, **(H)** Resistin), and BAT/BET markers (**(I)** EBF2, **(J)** TMEM26) in undifferentiated and differentiated cCPC (n≥4 independent cell culture experiments. Statistically significant differences between the groups are represented as mean±SD, mean of mRNA expression levels of undifferentiated cCPC were set at 100%, **P<0.01 vs. control, *P<0.05 vs. control, one sample *t*-test).

**S6 Fig. EPO supplementation prevents enhanced adipocyte formation in cultivated male CKO-CPC.**

**(A, B)** Bar graphs summarize ZFP423 mRNA levels in isolated male CPC incubated with rmEPO (10 ng/ml) for **(A)** 48 h or **(B)** 4 weeks after isolation (**(A)** : n=3 cell isolations, each isolation consists of 8-12 mice per genotype, **(B)** : n=5 independent isolations, each isolation consists of 10-12 animals per genotype). **(C)** Oil Red O staining visualizes adipocytes in CKO-CPC cultures after 4 weeks of cultivation with or without the addition of rmEPO (10 ng/ml), scale bars: 50 μm. **(D)** Bar graph summarizing adipocyte counts (n=5 independent isolations, each isolation consists of 10-12 animals per genotype). **(E, F)** qRT-PCR visualizes mRNA levels of the adipocyte markers **(E)** CEBPA and **(F)** FABP4 after 4 weeks of cultivation (n=3 independent cell isolations, each isolation consists of 10-12 animals per genotype). **(G)** Bar graphs summarize EZH2 mRNA levels in isolated CPC incubated with rmEPO (10 ng/ml) for 48 h (n=3 cell isolations, each isolation consists of 8-12 mice per genotype). **(A, B, D-G)** Data are presented as mean±SD, mean of WT PBS was set at 100 %, *P<0.05, **P<0.01 vs. WT PBS, ^#^P<0.05, ^##^P<0.01 vs. CKO PBS, two-way ANOVA with Bonferroni’s multiple comparison test.

